# novoRNABreak: local assembly for novel splice junction and fusion transcript detection from RNA-seq data

**DOI:** 10.1101/2022.12.16.520791

**Authors:** Yukun Tan, Vakul Mohanty, Shaoheng Liang, Jun Ma, Kun Hee Kim, Marc Jan Bonder, Xinghua Shi, Charles Lee, Human Genome Structural Variation Consortium, Zechen Chong, Ken Chen

## Abstract

We present novoRNABreak, a unified framework for cancer specific novel splice junction and fusion transcript detection in RNA-seq data obtained from human cancer samples. novoRNABreak is based on a local assembly model, which offers a tradeoff between the alignment-based and de novo whole transcriptome assembly (WTA) approaches, namely, being more sensitive in assembling novel junctions that cannot be directly aligned, and more efficient due to the strategy that focuses on junctions rather than full-length transcripts. The performance of novoRNABreak is demonstrated by a comprehensive set of experiments using synthetic data generated based on genome reference, as well as real RNA-seq data from breast cancer and prostate cancer samples. The results show that novoRNABreak can detect novel splice junctions and fusion transcripts efficiently with high sensitivity and reasonable specificity.

## INTRODUCTION

Splice junctions are conserved structures in eukaryotic genome that are recognized by RNA splicing machinery. Alternative splicing is one of the reasons for the production of many different transcripts (isoforms) from the same genetic locus. Dysregulation of RNA splicing has been found to be associated with many human diseases (1, 2), and established as one of the hallmarks of cancer (3). Fusion transcripts, resulting from gene fusion have been reported to be the driver mutations in neoplasia (4), including TMPRSS2-ERG in prostate cancer (5), BCR-ABL1 in chronic myeloid leukemia (6), and EML4-ALK in non-small-cell lung cancer (7). Thus, the identification of the junctions that provides valuable insights into alternative splicing and gene fusion events is biologically important and can potentially apply to cancer diagnosis, prognosis, and therapy (8).

With the advancement of next-generation sequencing (NGS) technologies, rapid and cheap genome-wide transcriptome analysis makes comprehensive detection of junctions possible. However, most of the available tools for junction detection primarily rely on approaches which directly align paired-end short reads to the genomic reference and identify the junctions from discordant read pairs, such as TopHat (9), Bellerophontes (10), Chimerascan (11), TumorFusions (12), INTEGRATE (13). Although computationally efficient, alignment-based approaches are fundamentally limited in detecting sequences that are substantially different from the reference, as such are most likely containing novel junctions due to challenges in accurately splitting and aligning short fragments. On the other hand, de novo whole transcriptome assembly (WTA) approaches, such as MINTIE (14), KisSplice (15), and TAP (16), which attempt to assemble all reads into a single consensus transcriptome, are computationally intensive and require high sequence coverage to achieve high sensitivity in assembling junctions. In the paper, we developed a new local-assembly-based pipeline to overcome those drawbacks by offering a tradeoff between the alignment-based and the de novo whole transcriptome assembly (WTA) approaches.

In this study, we proposed a local assembly-based framework, called novoRNABreak, which modifies our well-attested genomic structural variation breakpoint assembly tool novoBreak (17) to assemble novel junctions in RNA-seq data. It is a unified framework for novel splice junctions and fusion transcripts detection, which can identify the novel splice junctions and fusion transcript events according to the location of the splicing (one gene or two separate genes). The schematic diagrams of those events are shown in Figure 1 and Figure 2. With our k-mer guided local assemble model, our tool can fully use the unaligned sequences and is more sensitive in detecting clustered junctions formed by multiple short exons. As we will show in our experiment, more than 90% of the unmapped reads can be aligned confidently as assembled contigs after using our framework, indicating superior sensitivity of our tool in assembling structurally altered sequences in RNA-seq data. In addition, we argue that many alignment-based approaches, e.g., TopHat (9), SpliceMap (18), etc, are unable to find junctions when the reads span more than two small exons since they will only cut reads into two halves to do the alignment. Although SOAPsplice (19) proposed a strategy that splits a read into two segments of equal size for reads between 50bp to 100bp, and into multiple segments of 50 nucleotides for reads longer than 100bp, it didn’t solve the root of the problem and may cause other issues, e.g. the performance could get worse with reads longer than 101 bp, which was pointed out by (20).

**Figure 1.**
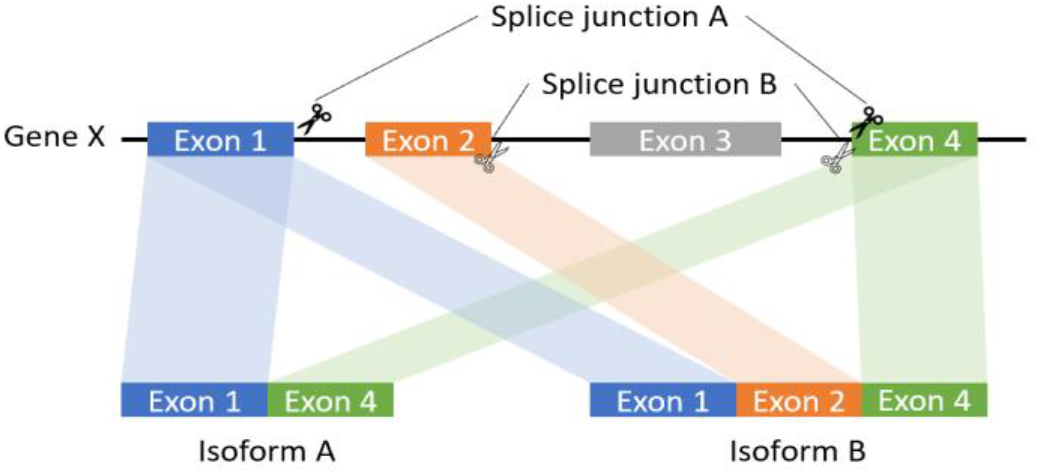
The schematic diagram of splice junction: sequences to aid in the process of removing introns by the RNA splicing machinery of one gene.

**Figure 2.**
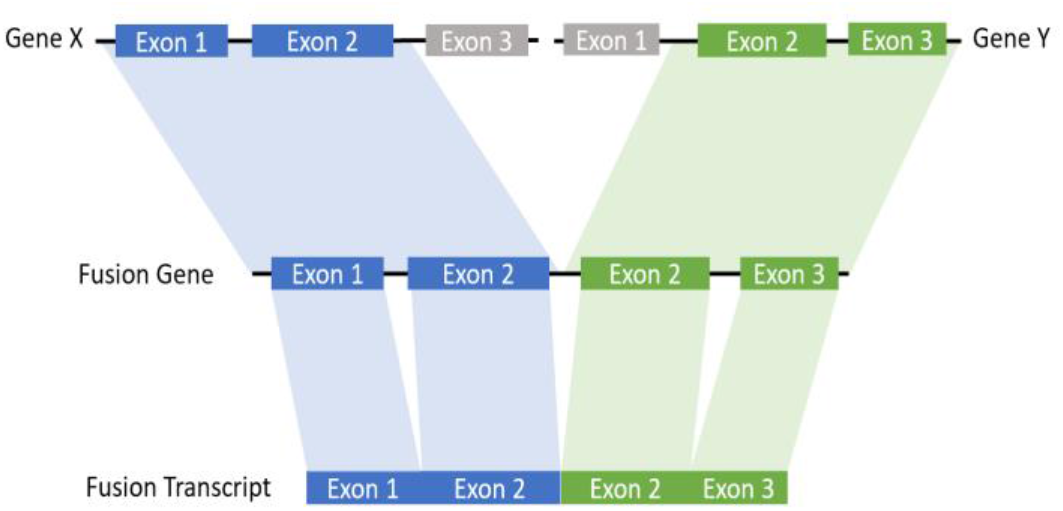
The schematic diagram of fusion transcript: a hybrid RNA is composed of transcripts of two separate genes.

The performance of novoRNABreak is demonstrated by a comprehensive set of experiments, including synthetic data generated from the genome reference, as well as real RNA-seq data from breast cancer, and The Cancer Genome Atlas (TCGA) prostate (PRAD) cancer samples. Results on the published real datasets followed by the experimental validated groundtruth show that our tool achieves higher sensitivity, especially in the cases where reads formed by multiple small structures, e.g., exons < 20bp, and reads cannot directly align to the reference.

## MATERIAL AND METHODS

### Alignment strategy

novoRNABreak, which modifies our well-attested genomic structural variation breakpoint assembly tool novoBreak, assembles novel junctions from RNA-seq data. Unlike many alignment-based or WTA approach methods in the literature, novoRNABreak consists of 4 steps shown in the Figure 3: First, RNA-seq reads and reference sequences will be decomposed into k-mers. We default to 31 as the k-mer size to achieve a balanced performance (17) and pick standard transcriptome databases such as NCBI RefSeq (21), Ensembl (22) and GENCODE (23) as the reference. Second, novel splice junction k-mers, which are absent in either the reference transcriptome or the normal samples but unique in the tumor RNA-seq reads, will be identified. Third, reads containing novel k-mers will be partitioned into clusters and assembled into sequences contigs using SSAKE (24), meaning that each of the contig contains at least one novel junction. Finally, the assembled contigs, which are now considerably longer than raw reads, are aligned against the Human GRCh38 genomic reference from Ensembl using Burrows-Wheeler Aligner (BWA) (25). Based on the alignment, the preliminary candidates of the junctions can be detected.

**Figure 3.**
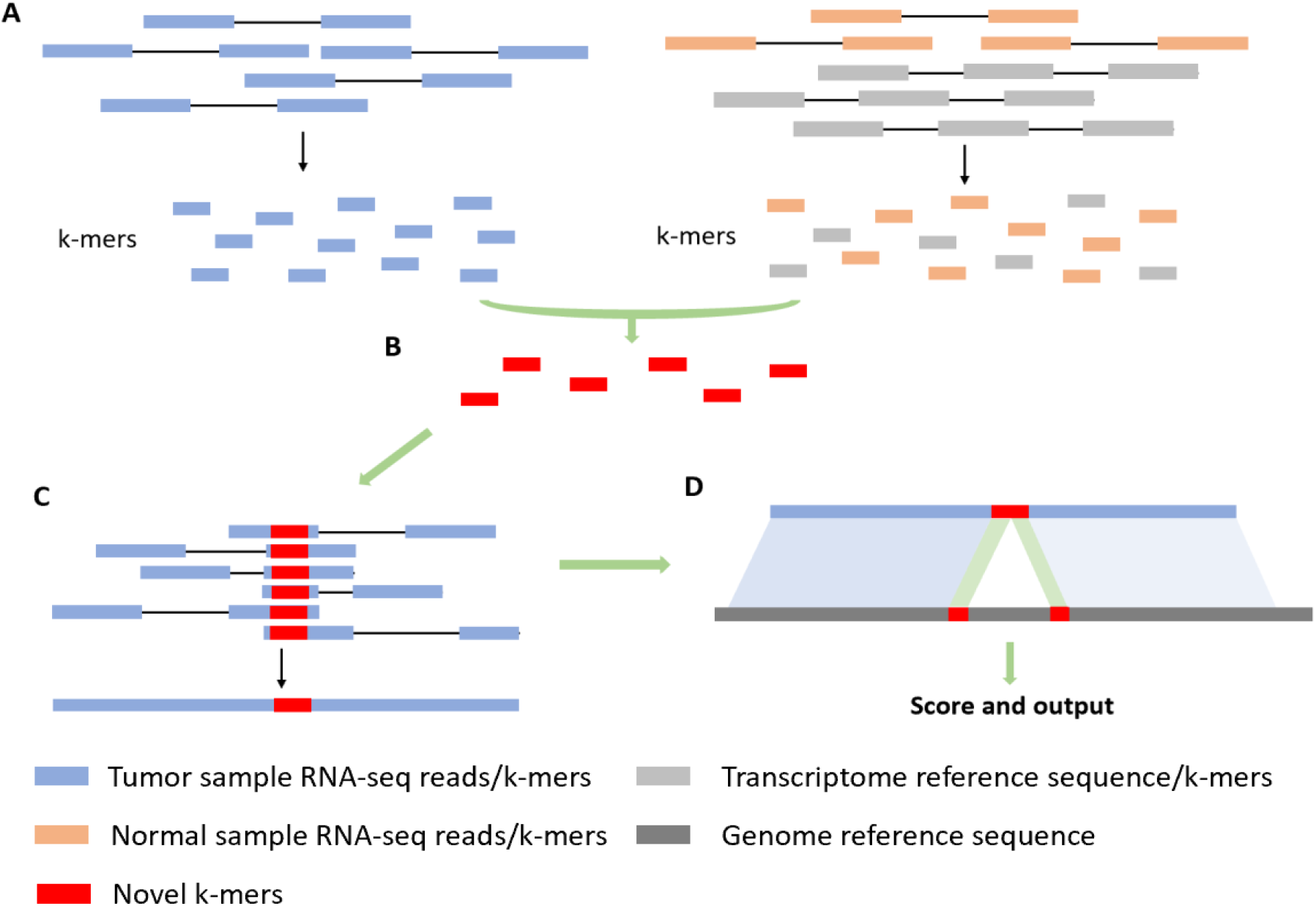
Alignment strategy. (A) Decompose RNA-seq reads and reference into k-mers. (B) Identify novel k-mers from tumor samples compared to normal samples and reference. (C) Partition reads containing novel k-mers into clusters and assemble into contigs. (D) Align against the genomic reference.

**Figure 4.**
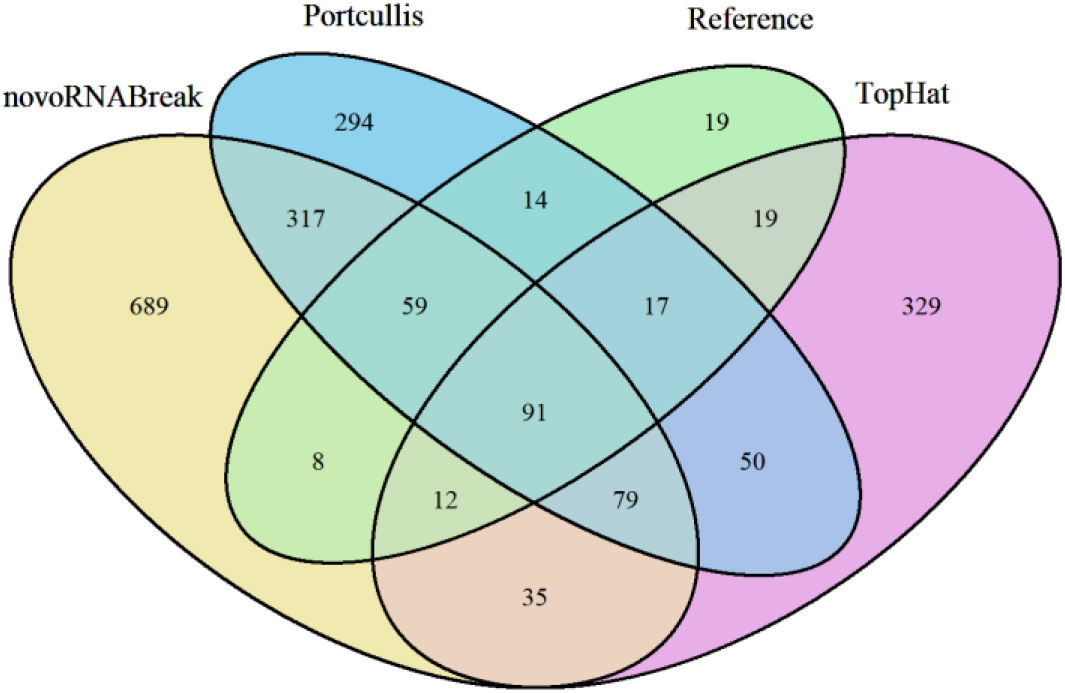
A four-way Venn diagram showing levels of agreement between novoRNABreak, Portcullis, TopHat, and the truth set (Reference) with 76 bp simulated reads.

### Filtering strategy

Our framework has one general filter set for both the splicing junction mode and the fusion transcript mode. There are also two specific filter sets for these two modes, respectively.

The general filter includes: (1) PCR-Artifact filter: It identifies and removes all duplicated reads introduced by the polymerase chain reaction (PCR) amplification process, e.g., Picard tool from Broad Institute (26). (2) Anchor length filter: Anchor length is the number of nucleotides overlapping each side of the break point and it can provide assurance of quality by removing all the junction-spanning reads having the anchor length lower than a threshold, e.g., 10bp. (3) Quality-Based filter: It uses the mapping quality parameter in the sam/bam file to discard the candidates with the mapping quality lower than a threshold. (4) Junction-Spanning reads filter: It considers the number of reads supporting the detected junctions and deletes the candidates with the number of supporting reads lower than a threshold, e.g., 3 reads, except when the contig is assembled by many short reads (at least 5) and has a high mapping quality (at least 60) at the same time. Note, the filter (1) and filter (4) are based on the actual mapped reads, and the filter (2) and (3) are based on the ensembled contigs.

The fusion transcript specific filters are: (1) Read-Through transcripts filter. It removes the RNA molecules formed by exons of adjacent genes, usually generated by the RNA-polymerase failing the recognition of the gene end. (2) Homology-Based filter: It is designed to remove the artifacts that are resulting from misalignment of read sequences due to polymorphisms and homology (27–29), e.g., HLA genes. (3) Ribosomal RNA-Based filter: It will remove highly expressed genes that are unlikely to be involved in fusions, such as ribosomal RNA (27, 29).

The splicing junction specific filters are: (1) Junction length filter: We limit the length of the junction in the range of 50 to 50,000 bp as this range covers most of the known intron size in eukaryote (19). (2) Canonical splice filter: The canonical splice sites are those with “GT” at the donor site, and “AG” at the acceptor site (also with “GC-AG” and “AT-AC”), which covers more than 99% of introns. We further filter our candidates by only allowing the form “GT-AG”, “GC-AG”, or “AT-AC” as the boundary of an intron, and “GT-AG” will be given the highest priority, followed by “GC-AG” and “AT-AC” boundaries (30). Note: genes are annotated by ANNOVAR (31) and junctions are annotated by RegTools (32).

## RESULTS

### Synthetic dataset

We simulated a dataset of 5,000,000 reads of length 76 bp using the R package ASimulatoR (33) and using human chromosome 21 as the reference. Here, we compare our novoRNABreak with Tophat (9) and Portcullis (20). A four-way Venn diagram shows the levels of agreement between the comparison tools and the truth set. Since our tool can detect the canonical novel junctions directly, we tweaked the truth set and other tools’ results to only keep the novel junctions. We observed that our tool can detect a competitive high number of canonical novel junctions, although has high false positive candidates. Accessing the dataset, we noticed that more than 99.9% simulated reads can be mapped to the reference directly, which is of great benefit to alignment-based approaches. Also, the scenario of reads spanning more than two small exons is rare and no novel junctions depending on that, which means this idealized data cannot demonstrate the advantages of our method but increase the number of misjudgments due to our model’s complexity. To demonstrate our ability to find junctions when the reads span more than two small exons, we generated a small dataset with still 76 bp length of the reads. We picked the exons of DOCK7 to design the required GTF input file as it has many small exons (length less than 25bp), meaning that the reads are more likely to span multiple junctions. The result is as we expected that only our tool can detect those junctions. We will show in the next section that this scenario persists on real datasets.

### Real dataset

We first evaluated the performance of the novel splicing junction detection mode of our tool by using the PRAD samples from TCGA. There are 499 tumor samples and 53 non-neoplastic samples. As explained in (12), non-neoplastic samples in TCGA are frequently obtained through tissue biopsy adjacent to the location of the cancer which have the risk of being contaminated with tumor cells. We identified 7 out of 53 non-neoplastic samples as true nor mal using unsupervised clustering. With those normal samples, our tool can directly deliver the cancer specific novel canonical junctions. By removing the junctions which output by Portcullis and TopHat, we detected some consensus cancer specific novel canonical junctions that are only found by our method. We also noticed that many genes where we detected the novel junctions are related to the prostate cancer, e.g., RANBP3L (34), NAAA (35), PYROXD2 (36), etc. Taking one of our detections, the junction from SYT17, as an example. The junction we detected is from 19172777 to 19173444 on chromosome 16. The sashimi plots in Figure 5, created by (37), depicted that novel tumor-specific splice junction. As shown in Figure 5, it is a novel combination of the known donor and acceptor. The donor relates to a small exon (19172760-19172777) from the transcript ENST00000568526, etc, with only 17 bp, and the acceptor position relates to another exon (19173444-19173578) from the transcript ENST00000568115. The original acceptor from ENST00000568526, etc, beginning at 19173430 has been truncated. We can see that those exons are small enough to have a high possibility to be covered by a single read which are difficult for most of the alignment-based methods. Next, we checked the effect of the novel junction on open reading frames (ORFs) by using the online tool from NCBI (https://www.ncbi.nlm.nih.gov/orffinder/). The comparison between Figure 6 and Figure 7 shows that the 14 bp truncation will change the position of the start codon and the end codon of all 6 possible reading frames, which may further affect the translation of the protein.

**Figure 5.**
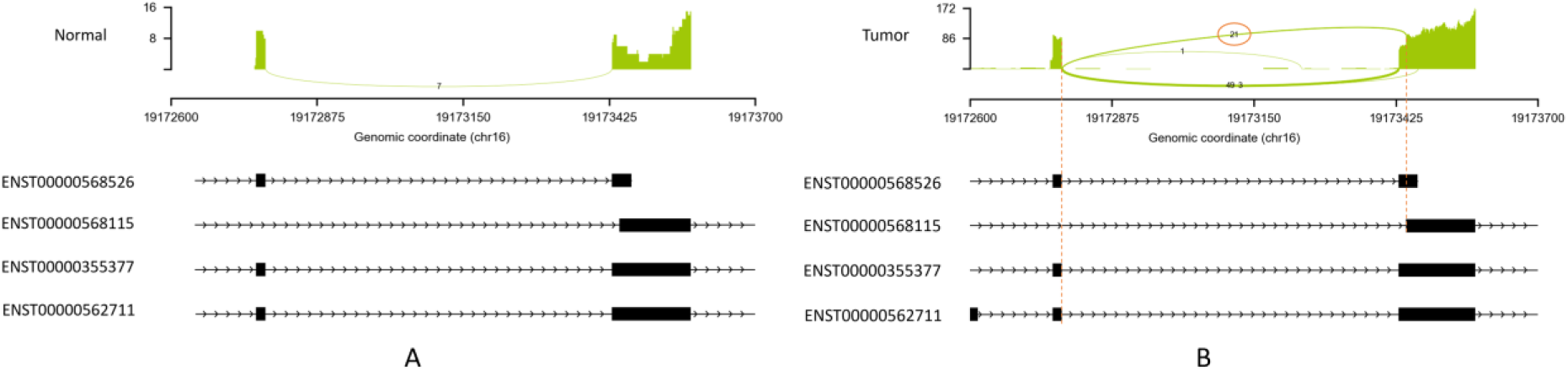
Sashimi plots depicting tumor specific splice junction. (A) Splice junction on a normal sample. (B) Splice junction on a prostate cancer sample.

**Figure 6.**
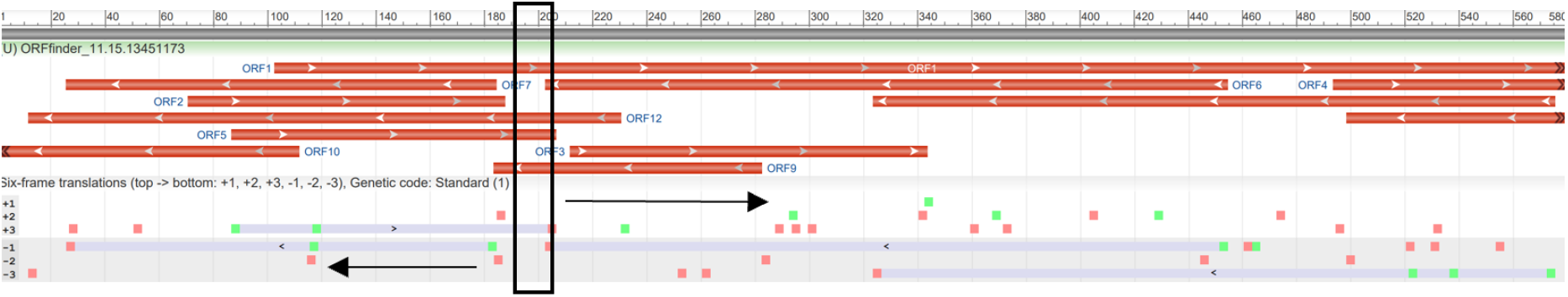
Six possible reading frames of the original transcript ENST00000355377. The black frame is the 14 bp truncated part.

**Figure 7.**
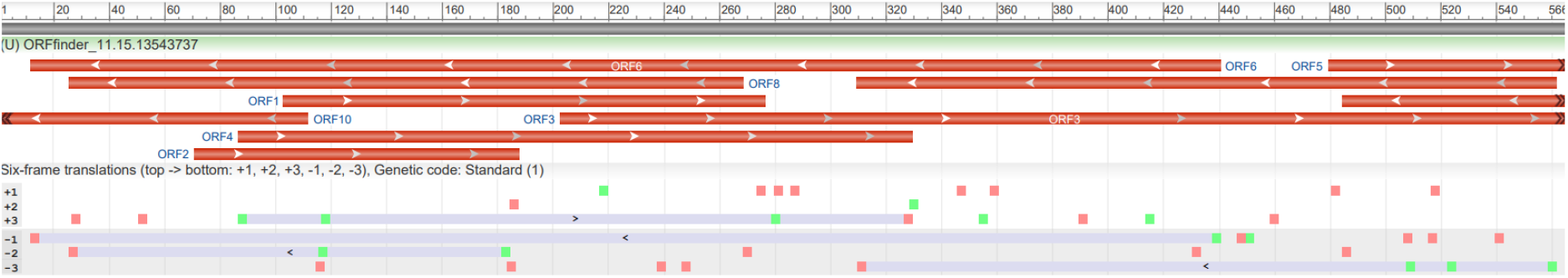
Six possible reading frames of the novel transcript.

Furthermore, we evaluated the performance of the fusion transcripts detection mode of our tool using several real datasets based on human genome reference sequence, including breast cancer and TCGA PRAD samples. The breast cancer dataset in this study consists 4 cell lines (BT-474, SKBR-3, KPL-4, and MCF-7) which can be downloaded from NCBI Sequence Read Archive (SRA) with accession number SRP003186 (38). There are total 26 experimentally verified fusion events for breast cancer cell lines (The fusion CSE1L-ENSG00000236127 was removed from the list due to the deprecation of ENSG00000236127) (39). The comparison results are shown in Figure 8, where the outcomes of other methods (11, 40–48) are picked from the review paper (49). We can see that our tool detects the most of validated fusion transcript in total, although not the best in every cell line. We can reach a high sensitivity because our method can fully utilize the unmapped data. There are the total of 198,714,026 reads from those 4 cell lines, of which 7,341,176 reads are unmapped (3.7%). By using those unmapped reads only, we assembled 106,574 high-quality contigs, in which 8 true fusion transcripts can be identified and 5 of them passed all the filters (high quality). More importantly, 2 of them have no support from the mapped short reads, meaning that those 2 would theoretically be missed by the alignment-based methods.

**Figure 8.**
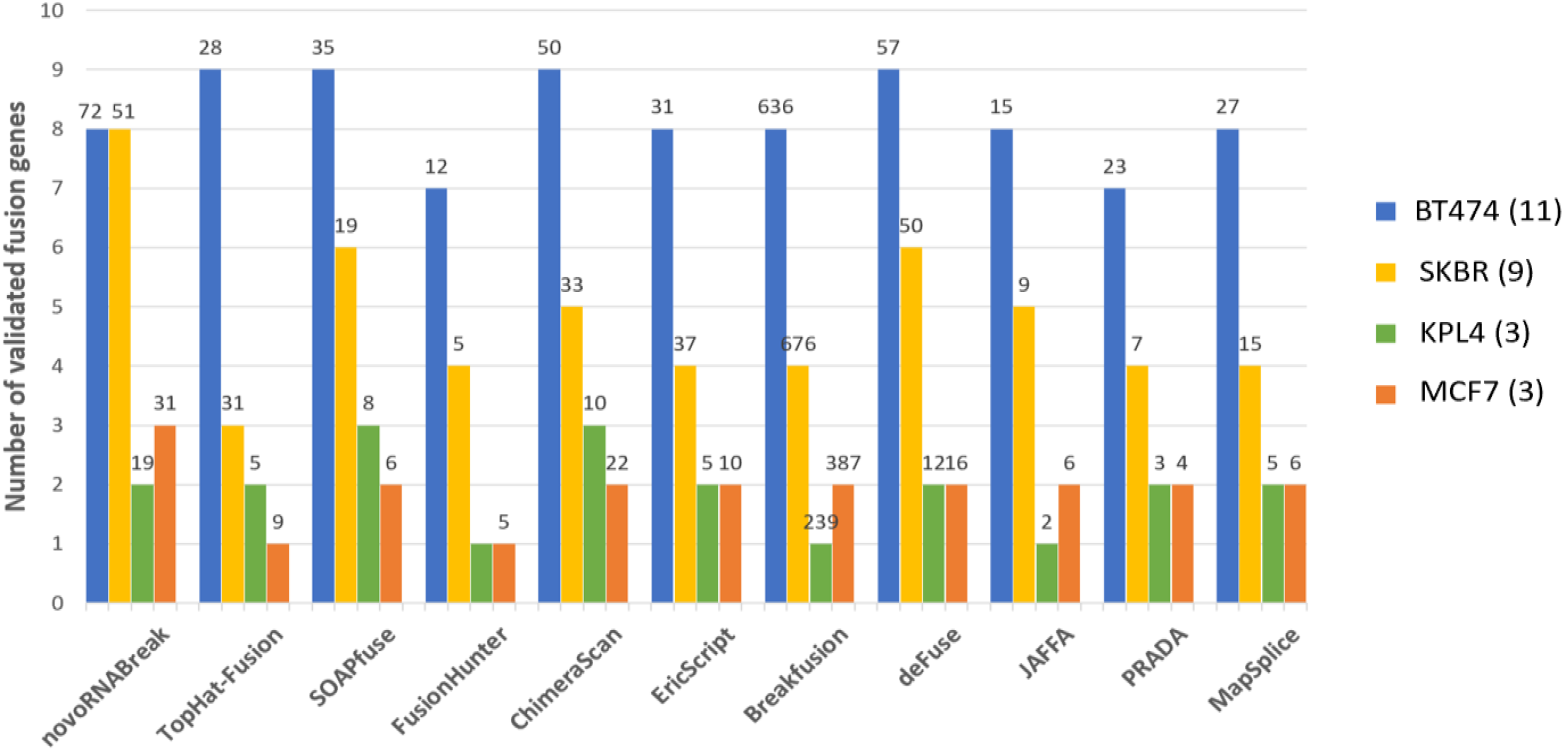
Fusion transcript detection results for the real breast cancer data set. The y-axis bars show the number of true detected positives (benchmarks). The total number of fusion detections are shown on the top of the bars.

In addition, we applied our framework to the TCGA PRAD samples, the same dataset as we used to evaluate our novel splicing junction mode. We compared the results with INTEGRATE (13) and TumorFusions (12) which are also applied to TCGA PRAD samples. By using the prostate cancer fusion genes list from the review paper (50), we extracted the consensus fusion events and show in Table 1. As we expected, we can recover more fusion events. The full list of the detected fusion events from our tool is included in the supplementary material.

**Table 1.** Fusion events comparison between novoRNABreak, INTEGRATE, and TumorFusions.

| Fusion genes | novoBreak-rna | INTEGRATE | TumorFusions |
| --- | --- | --- | --- |
| TMPRSS2-ERG | ✓ | ✓ | ✓ |
| SLC45A3-ERG | ✓ | ✓ | ✓ |
| TMPRSS2-ETV4 | ✓ | ✓ | ✓ |
| TMPRSS2-ETV1 | ✓ | ✓ | ✓ |
| SLC45A3-ETV1 | ✓ | ✓ | ✓ |
| KLK2-FGFR2 | ✓ |  | ✓ |
| TMPRSS2-ETV5 | ✓ |  | ✓ |
| NDRG1-ERG | ✓ | ✓ |  |
| ACER3-B3GNT6 | ✓ |  |  |
| KLK2-ETV1 | ✓ |  |  |
| ACPP-SKIL | ✓ |  |  |

## DISCUSSION

Here we present a unified framework for identifying tumor specific novel canonical splicing junctions and novel fusion transcripts from RNA-seq data. Our results suggest that our tool has a better performance by fully utilizing unmapped reads and by sensitively identifying the junctions when reads spanning two or more small exons. Furthermore, the novel events detected from our method will improve our understanding of cancer mechanisms and facilitate discovery of new targets and development of RNA-based therapies.

## DATA AVAILABILITY

The source code for novoRNABreak is freely available on GitHub (https://github.com/KChen-lab/novoRNABreak).

## SUPPLEMENTARY DATA

Supplementary Data are available at NAR online.

## FUNDING

This project has been made possible in part by the National Cancer Institute Informatics Technology for Cancer Research [U01CA247760 to KC], the Cancer Prevention & Research Institute of Texas [RP180248 to KC], the National Cancer Institute Cancer Center Support [P30 CA016672 to PP], the National Institute of General Medical Sciences [1R35GM138212 to ZC] and the National Human Genome Research Institute [U24-HG-007497 to CL].

## Notes

### Competing Interest Statement

The authors have declared no competing interest.

